# The Impact of Reference Genome Divergence on Ancient DNA Damage Detection in Metagenomic Contexts

**DOI:** 10.1101/2025.07.16.665190

**Authors:** Benjamin Guinet, Bilal Muhammad Sharif, Tom van der Valk

## Abstract

The reliability of ancient DNA (aDNA) authentication depends on detecting characteristic damage patterns, particularly cytosine deamination at fragment ends. However, in ancient metagenomic studies, sequence divergence between aDNA reads and available reference genomes may obscure such damage signals. We systematically evaluated how reference genome divergence, read count, read length, and damage levels affect aDNA damage profiles using both empirical datasets and controlled simulations. Using ancient Yersinia pestis and Hepatitis B virus data, we show that mapping to divergent reference genomes significantly reduces the detectability and intensity of characteristic damage patterns, particularly at low read counts. Simulations further revealed that reference genome identity is the strongest predictor of damage intensity, while read count primarily influences damage stochasticity. We introduce a correction matrix that adjusts C-to-T damage profiles for reference divergence, improving damage signal recovery. Our findings highlight methodological considerations for authenticating aDNA in metagenomic contexts, particularly when closely related reference genomes are unavailable.

## INTRODUCTION

Ancient DNA can offer valuable insights into past environments; however, reliably distinguishing authentic ancient molecules from modern contaminants is critical for accurate reconstructions. After an organism dies, DNA repair mechanisms stop, and the DNA gradually fragments, forming single-stranded overhangs at the molecule ends. These overhangs expose cytosines to environmental damage, particularly deamination, which converts cytosine to uracil. During sequencing library amplification, these uracils are misread as thymines, resulting in characteristic damage patterns, most notably an increase in cytosine-to-thymine (C-to-T) substitutions at the ends of DNA fragments (Briggs et al. 2007) (Bokelmann, Glocke, and Meyer 2020). A key authentication method in ancient DNA is thus to show the presence of elevated C to T deamination patterns (and the complementary G to A) at the sequence read ends, which when plotted show a characteristic “smiley” pattern.

However, aside from biological variation, factors such as genome coverage, and the amount of damage present in the DNA fragments have been shown to influence ancient DNA authentication (Borry et al. 2021). Typically, ancient DNA damage patterns are measured by aligning reads to a reference genome from the same or a closely related species. Yet, with the rise of microbial and viral inference studies from ancient materials such as dental calculus and sediments, researchers commonly encounter DNA from organisms that lack a reference genome (Kjær et al. 2022; Fernandez-Guerra et al. 2023). In these cases, reads are aligned to a phylogenetically close relative with a reference available in genomic databases, often resulting in substantial sequence divergence between the ancients reads and the used reference genome. Moreover, the analysis of deep-time sequence data from microorganisms over a million years old can result in substantial sequence divergence even when a reference for the modern species is available (Guinet et al. 2025, Kjaer et al 2022). This large number of biological substitutions to the reference, complicates the estimation of C-to-T damage rates, as the distinct damage signal can become overwhelmed by the high overall substitution rate. Furthermore, ancient metagenomic studies often recover only a small number of sequence reads for the species of interest, further reducing the statistical power to accurately estimate damage rates. Finally, aligning the typically short ancient DNA reads to a divergent reference genome introduces mapping biases, which become more pronounced with increasing reference divergence (Dolenz et al. 2024). The combination of these factors may prevent the detection of characteristic ancient DNA damage patterns, even if the reads originate from authentic ancient DNA. With the increasing number of ancient metagenomic studies, a systematic assessment of how reference genome divergence, read count and read length quantitatively affect ancient DNA damage estimates enhances our understanding of expected damage patterns across datasets and supports more robust downstream authentication methods. In this study, we employ an extensive set of simulated and empirical microbial and viral paleogenomic datasets to systematically investigate the impact of key factors, including read length, divergence to reference genome, damage levels and reads depth, on the characterization of aDNA damage patterns.

## RESULTS

To assess how the aforementioned variables influence DNA damage profiles, we first analyzed ancient sequence data from two organisms: 222,117,414 sequence reads previously obtained from a 1360 year old *Yersinia pestis (Namouchi et al. 2018)* and 16,454 sequence reads from a 2440 year old *Hepatitis B virus* genotype B (HBV-B) (Sun et al. 2024). We aligned this data to their respective modern reference genomes, as well as a set of references from divergent but related species/strains. While both these samples show a clear signal of ancient DNA when aligned to their respective references, subsampling to lower read numbers introduces high stochasticity (**Fig. 1** and **Fig. 2**). Nonetheless, when mapping *Y. pestis* and HBV-B to their respective reference genomes, even a modest number of reads (n = 500) was sufficient to produce the characteristic “smiley” damage profile, with a clear increase of C-to-T mismatches at the ends of reads (**Fig. 1** and **Fig. 2**). Even at only 100 reads, DNA damage can be detected, albeit with high stochasticity (**Fig. 1**). Conversely, when mapping the sample reads to distant reference assemblies such as *Y. pestis* mapped to *Escherichia coli* (sequence divergence = 6.3%) or HBV-B to Woolly Monkey Hepatitis B virus (sequence divergence = 7.9%) a strong effect of reference genome divergence on damage estimates is detected. In both cases, 100 reads are insufficient to detect a damage pattern, and even with 500 aligned reads, the damage signal is highly stochastic and difficult to discern from random noise (**Fig. 1** and **Fig. 2**).

**Figure 1.**
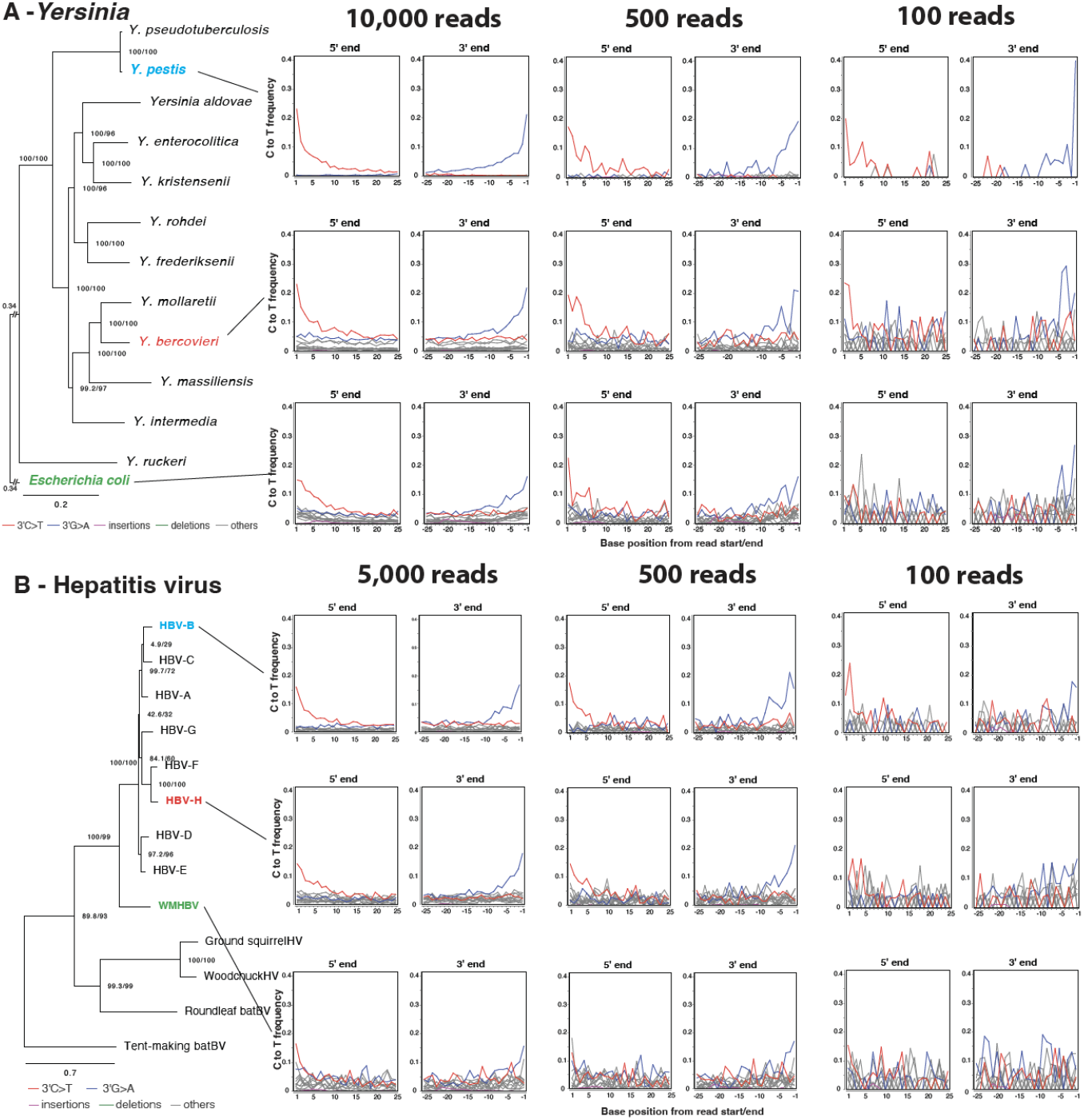
**A - Observed damage plots for ancient *Yersinia pestis* reads mapped to increasingly divergent genome assemblies with different numbers of reads.** Top row (Blue taxa) represents reads mapped to the reference assembly *Yersinia pestis*, middle row (red) to the distantly related *Yersinia bercovieri* assembly and bottom row (green) to *Escherichia coli*. **B - Observed damage plots of *Hepatitis B virus* reads mapped to increasingly divergent genome assemblies with different numbers of reads**. Top row (Blue taxa) represents reads mapped to the reference assembly Hepatitis B virus, middle row (red) to another distantly related Hepatitis B virus from the group H and bottom row (green) to the Woolly Monkey Hepatitis B virus. Phylogenies were built with Iqtree2 based on the alignment of the assemblies of the reference genome of each taxa using Sibeliaz. Confidence scores (aLRT%/ultra-bootstrap support%) are shown at each node. Branch-length scale is shown at the bottom of the phylogeny. Damage plots were made using DamageProfiler.

**Figure 2.**
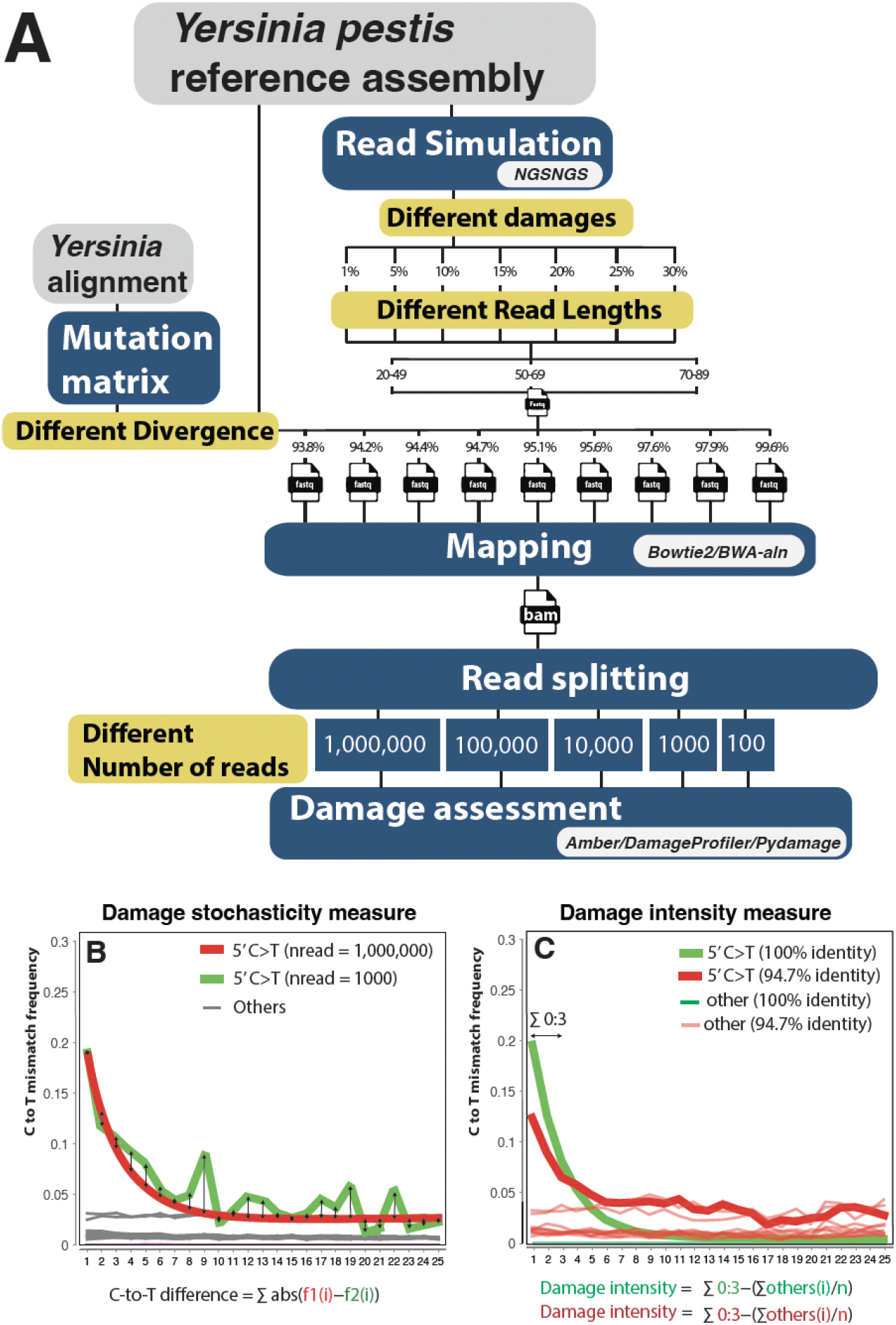
Schematic overview of the simulation pipeline and metrics used to quantify damage profile stochasticity and intensity. **(A)** - For the simulated data, we first generate 1,000,000 reads from the *Yersinia pestis* reference assembly using NGSNGS, applying different levels of damage ranging from 1% to 30%. This percentage represents the maximum C-to-T mismatch rate at the first position of the reads. Additionally, we simulate reads with varying lengths, uniformly distributed between 20–49 bp, 50-69bp and 70–89 bp. To mimic genome divergence, mutations are introduced into the reads. This is achieved by calculating a mutation matrix derived from sequence alignments between several *Yersinia* species genomes. The simulated divergence ranges from 93.8% to 99.6% (including the simulated DNA damage mutations). The simulated reads are then mapped to the *Yersinia pestis* reference genome using both Bowtie2 and BWA-aln. To estimate the effect of read depth, we downsampled the mapped reads in the BAM files, ranging from 100 to 1,000,000 reads. Finally, DNA damage assessment is performed using Amber, DamageProfiler, and PyDamage, three complementary tools that together provide a comprehensive analysis of DNA damage. (**B)** The red curve illustrates the observed damage profile when up to 1 million reads are mapped to a given reference genome. The green curve represents a profile obtained under different parameters and with fewer mapped reads. The C-to-T difference metric is calculated as the sum of absolute differences at each position in the damage profile. Higher values indicate greater deviation of the green curve from the red curve, reflecting increased stochasticity. **(C)** The red and green curves represent distinct damage patterns, with the green curve corresponding to reads mapped to a distant reference genome with 94.7% identity, and the red curve to reads mapped to the identical reference genome (100% identity). To compare damage intensity at the read edges, we sum the C to T mismatch frequencies at the first three positions of each profile. To account for the overall higher mismatch rate in the red curve due to larger divergence to the reference, we apply a correction by subtracting the average mismatch frequency of all other mismatch types from the total sum.

To obtain further insights into the variables affecting these DNA damage signals, we designed a controlled *in silico* experiment to systematically assess the effects of read count and reference genome divergence. We also incorporated read length variables and different levels of DNA damage in the simulated sequence data as well as using different algorithms for read mapping. In these simulations, the *Y. pestis* modern reference (GCF_000222975.1) was used to simulate a wide range of parameters (see Materials and Methods). First, in 73.3% of the simulations we detected a significant difference (Wilcoxon test) between the damage curves produced by the most commonly used mapping tools for ancient DNA; Bowtie2 and BWA-aln. Along the entire read length, BWA-aln alignments showed a higher C-to-T mismatch frequency compared to Bowtie2 aligned reads (**FigS3**). We used the software Pydamage, which employs a likelihood ratio test to discriminate between ancient and modern contaminant reads to estimate the strength of aDNA damage signals among all the aligned reads. In 75.3% of the simulations a *P*-value < 0.05 was obtained for both the Bowtie2 and BWA-aln alignment. Additionally, 7.5% of the simulations showed a significant DNA damaged pattern in the Bowtie2 alignments but not the BWA-aln alignments, whereas the opposite scenario occurred in only 2.5% of the cases, finally in 14.7% of simulations, both analyses did not show any significant damage. These results may be explained by the observations that Bowtie2 was overall more stringent than BWA-aln. This is supported with BWA showing higher average mismatches than Bowtie2 (mean difference = 0.417, 95% CI: [0.371, 0.463], *p* < 0.001). This suggests that using Bowtie2 settings can enhance the recovery of expected damage patterns. For the remaining analysis of this study, we thus opted to use only Bowtie2 alignments.

We systematically evaluated the impact of read count, reference genome divergence, and damage level on the p-values obtained from Pydamage. Even though DNA damage was included in all our simulations, when testing for the authenticity of ancient DNA, the p-values for were often non-significant when the number of reads or the level of damage was low (**FigS8**), consistent with previous findings ((Borry et al. 2021; Briggs et al. 2010; van der Valk et al. 2021)). We demonstrate that as the reads diverge further from the used reference genome, the number of simulations yielding non-significant p-values increases markedly (**FigS8**).

To quantify the change in the level of estimated ancient DNA damage in a systematic way, we here established two metrics: **“damage level”** and **“damage stochasticity.” Damage stochasticity** assesses the level of noise in the C-to-T mismatch frequency curve as the average divergence from the simulation values for each base in the read (**Fig. 2A**), while **Damage level** measures the overall amount of DNA damage, calculated as the sum of the C-to-T substitution rate at the first and last 3 bases of the sequencing reads minus the average C-to-T substitution rate across all other bases in the read (**Fig. 2B)**.

### Damage intensity patterns are influenced by assembly divergence

When simulating 20% DNA damage (i.e. on average 20% of reads have damage at the first base), we observe a decrease in the detected damage as reads are mapped to increasingly divergent reference genomes. For example, with a reference genome diverging by 5%, the observed damage is reduced by 28% (i.e. from 20% to 14.4% at the first base) (**Fig. 3A**). When mapping to a highly divergent reference with 6.4% sequence divergence, the observed damage decreases by 59.5% (**Fig. 3A**). . he number of mapped reads had only a minor effect on the observed damage intensity, except when very few reads were mapped (e.g., *n* = 100). For example, mapping 5,000 reads resulted in a slight decrease in damage of just 0.34%, whereas mapping only 100 reads led to a more pronounced reduction of 1.48% (**Fig. 3B**). Finally, damage stochasticity seems more strongly influenced by the number of mapped reads; a low read count leads to greater variability in the damage profile (**Fig. 3B**). This effect is not observed when reads are mapped to distant reference assemblies (**Fig. 3A**).

**Figure 3.**
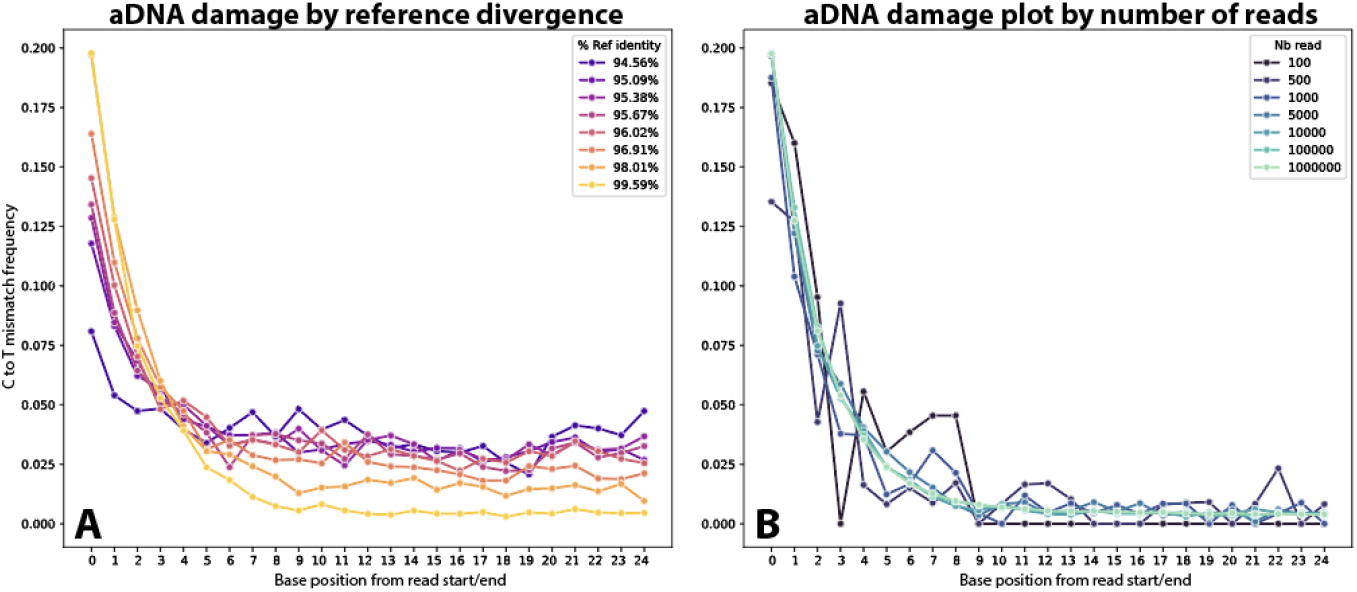
C to T mismatch frequency as a function of reference divergence and read numbers. **A** - Each curve shows the average C to T mismatch frequency at each position across 10,000 reads mapped the reference of increasingly lower sequence identity. **B** - Each curve represents the average C to T mismatch frequency at each position across all reads mapped to the same (*Yersinia pestis)* reference genome as the reads (i.e. 100% identity when excluding DNA damage). The dataset used in the A plot corresponds to 10,000 reads mapped. In both plots, simulated read lengths ranged from 50-59bp and simulated damage of 20%.

When comparing different levels of aDNA damage, ranging from 5% to 30%, we observe only a minor effect of read count on the damage estimates (**Fig4-A & FigS4**). However, damage estimates are highly influenced by the sequence divergence of the used reference genomes (**Fig4-A & FigS4**), with this effect being more pronounced at higher damage levels. At low damage levels, when reads exhibit only 5% damage, the difference in estimates across different reference genomes is negligible (**Fig4-A & FigS4**).

**Figure 4.**
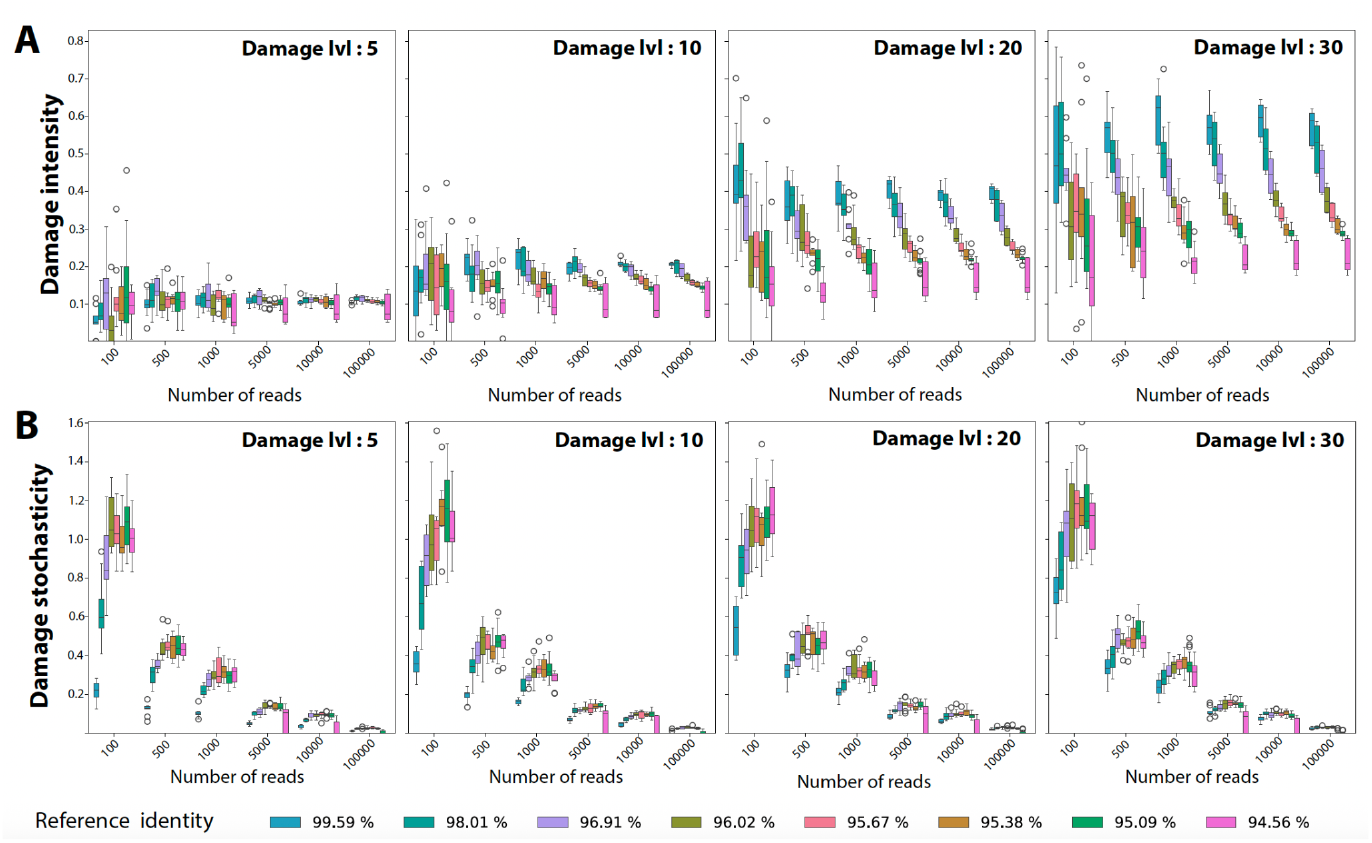
DNA damage estimates and stochasticity depending on the mapping conditions. **A -** Boxplots showing the distribution of damage levels across different read depths (nread) and reference genome percentage identities for varying damage levels. **B -** Boxplots showing the distribution of the damage stochasticity across different read depths (nread) and reference genome percentage identities for varying damage levels. DNA damage and stochasticity were calculated as shown in **Fig 2-A&B**.

To statistically test these observations further, we built a regression model to determine which of the variables, read length, damage level, reference divergence, and the number of reads explain the variation in damage intensity. We used the linear regression approach (*Damage intensity ∼ Damage_level + Nb_reads + Ref_percID + Read_length)*, implemented in Python using the statsmodels library and fitted the model to all the inferred values from the simulation data. The number of reads did not present a significant relationship with damage intensity (all p > 0.10) (**Table1**). The reference percentage identity variables showed the strongest correlation with damage intensity. For instance, the coefficients for 94%ID and 99.6%ID are respectively 0.043 and 0.129 (both p < 0.00001) (**Table1**), indicating that a close sequence identity between the reads and the reference genome is associated with a significant increase of the observed damage intensity. Finally, the read length variable exhibits mixed effects. Shorter reads such as 20-49 bp and 50-69 bp slightly reduced damage intensity (coefficient = -0.012 and -0.008 respectively, p < 0.00001)(**Table1**), while longer reads (70-89 bp) did no not have a significant effect, suggesting that read length plays a nuanced role in determining damage intensity, with short and long reads affecting it in different ways.

**Table 1.** Coefficients from ordinary least squares (OLS) regression predicting the effect of several predictors on the damage intensity. The table reports the estimated coefficient (coef), standard error (Std. Error), t-value, p-value, and the 95% confidence interval (CI) bounds.

|  | coef | std err | t | P>\|t\| | [0.025 | 0.975] |
| --- | --- | --- | --- | --- | --- | --- |
| Intercept | -0.0191 | 0.003 | -5.656 | 0.000 | -0.026 | -0.012 |
| Damage[1%] | 0.0116 | 0.004 | 2.929 | 0.003 | 0.004 | 0.019 |
| Damage[5%] | 0.0608 | 0.004 | 15.357 | 0.000 | 0.053 | 0.069 |
| Damage[10%] | 0.1208 | 0.004 | 30.329 | 0.000 | 0.113 | 0.129 |
| Damage[15%] | 0.1791 | 0.004 | 45.113 | 0.000 | 0.171 | 0.187 |
| Damage[20%] | 0.2316 | 0.004 | 58.306 | 0.000 | 0.224 | 0.239 |
| Damage[25%] | 0.2771 | 0.004 | 69.385 | 0.000 | 0.269 | 0.285 |
| Damage[30%] | 0.3317 | 0.004 | 83.037 | 0.000 | 0.324 | 0.339 |
| nread[500] | 0.0003 | 0.004 | 0.072 | 0.942 | -0.007 | 0.008 |
| nread[1000] | -0.0028 | 0.004 | -0.749 | 0.454 | -0.010 | 0.005 |
| nread[5000] | 0.0002 | 0.004 | 0.044 | 0.965 | -0.007 | 0.007 |
| nread[10000] | -0.0001 | 0.004 | -0.037 | 0.971 | -0.007 | 0.007 |
| nread[100000] | 0.0002 | 0.004 | 0.059 | 0.953 | -0.007 | 0.008 |
| nread[1000000] | 0.0002 | 0.004 | 0.062 | 0.950 | -0.007 | 0.008 |
| Ref_percID[94.0%] | 0.0434 | 0.004 | 9.967 | 0.000 | 0.035 | 0.052 |
| Ref_percID[94.19%] | 0.0356 | 0.004 | 8.193 | 0.000 | 0.027 | 0.044 |
| Ref_percID[94.45%] | 0.0454 | 0.004 | 10.441 | 0.000 | 0.037 | 0.054 |
| Ref_percID[94.85%] | 0.0582 | 0.004 | 13.380 | 0.000 | 0.050 | 0.067 |
| Ref_percID[95.29%] | 0.0683 | 0.004 | 15.689 | 0.000 | 0.060 | 0.077 |
| Ref_percID[96.53%] | 0.1029 | 0.004 | 25.984 | 0.000 | 0.095 | 0.111 |
| Ref_percID[97.79%] | 0.1242 | 0.004 | 28.551 | 0.000 | 0.116 | 0.133 |
| Ref_percID[99.57%] | 0.1296 | 0.004 | 29.776 | 0.000 | 0.121 | 0.138 |
| Read_Length[50-69bp] | -0.0079 | 0.002 | -4.189 | 0.000 | -0.012 | -0.004 |
| Read_Length[70-89bp] | 0.0012 | 0.001 | 1.138 | 0.255 | -0.001 | 0.003 |
| Read_Length[20-49bp] | -0.0124 | 0.002 | -6.573 | 0.000 | -0.016 | -0.009 |

### 2. stochasticity patterns are primarily influenced by the number of reads

Across all damage levels (from 5% to 30%) we observe a clear correlation between the number of mapped reads and the stochasticity of the damage curves. Specifically, greater stochasticity is observed with fewer mapped reads (**Fig. 4-B & FigS7**). This trend remains consistent across varying degrees of damage. To a lesser extent, we also observe a slight reduction in damage stochasticity when reads are mapped to closely related reference genomes. and this effect appears to be more pronounced with a small number of reads (**Fig. 4-B & FigS7**).

When statistically testing these observations (*Damage stochasticity ∼ Damage_level + Nb_reads + Ref_percID + Read_length)*), our analysis shows that damage levels are significantly associated with stochasticity of the damage curve. As the damage level increases, the observed difference in stochasticity becomes more pronounced. For example, at higher damage levels (such as 30%), the coefficient reaches 0.07 (p < 0.00001), while at lower damage levels (like 5%), the coefficient drops to 0.02 (p = 0.001) (**Table2**). The number of mapped reads is the main factor explaining damage stochasticity. As the number of reads increases, the damage stochasticity decreases. This is evident from the coefficients for different read categories, such as nread=1000 (-0.66, p < 0.00001) and nread=1,000,000 (-0.92, p < 0.00001)(**Table2**), showing that larger sequence datasets reduce the magnitude of the observed differences in substitutions. There is also a notable decrease in stochasticity as the reference identity increases. For example, at 99.6% identity, the coefficient is -0.15 (p < 0.00001)(**Table2**), indicating that higher alignment precision with the reference genome results in a smaller stochasticity. At lower identity levels (e.g., 94 %ID), the effect is positive, with a coefficient of 0.02 (p < 0.00001)(**Table2**), showing that lower reference identity corresponds to larger stochasticity. Read length also significantly impacts the damage stochasticity. Shorter reads (20-49 bp) increase stochasticity (coefficient = 0.24, p < 0.00001), whereas longer reads (70-89 bp) show a substantial decrease (coefficient = 0.14, p < 0.00001)(**Table2**).

**Table 2.**
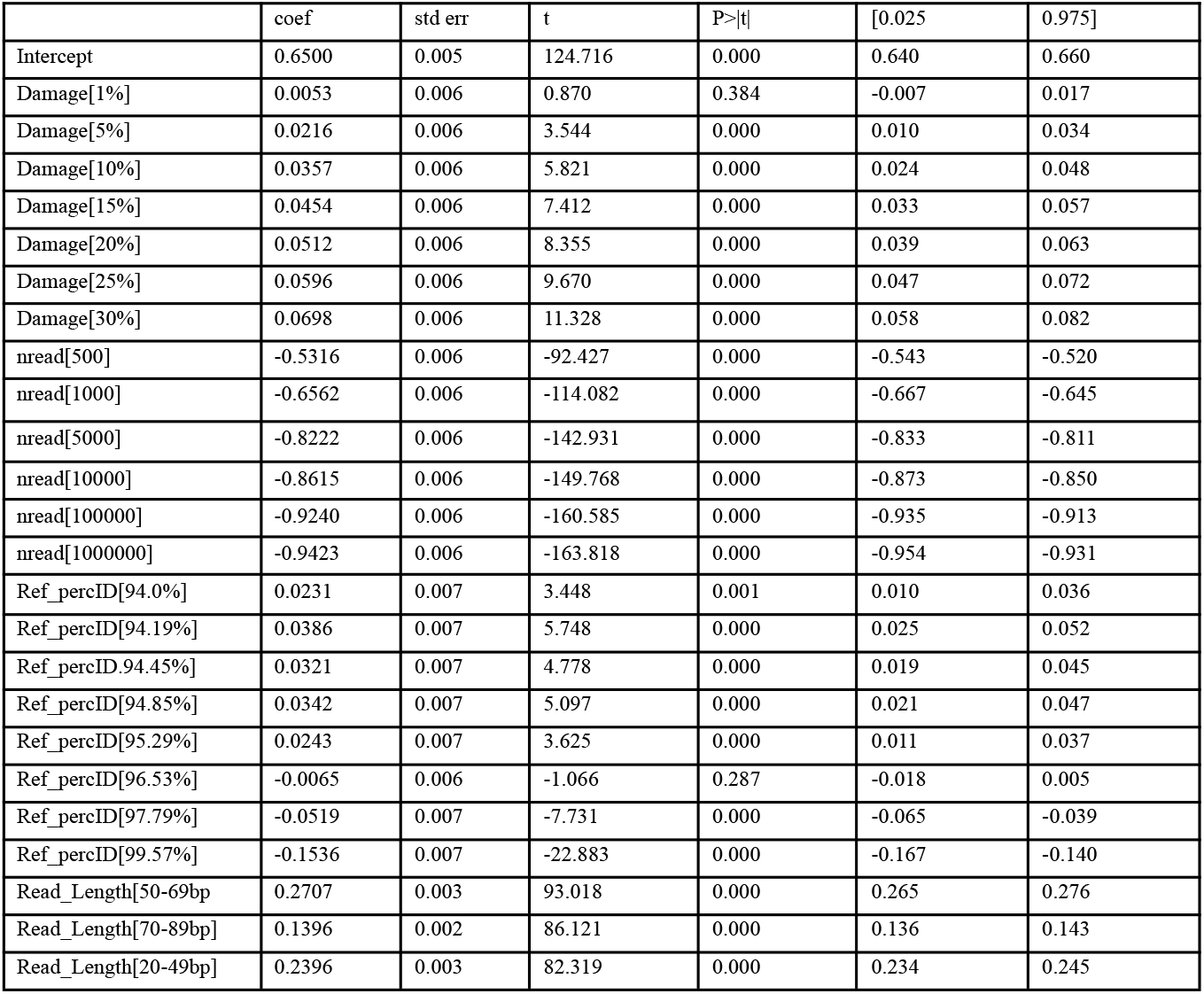
Coefficients from ordinary least squares (OLS) regression predicting the effect of several predictors on the damage stochasticity. The table reports the estimated coefficient (coef), standard error (Std. Error), t-value, p-value, and the 95% confidence interval (CI) bounds.

### 3. Correcting DNA damage estimates for reference genome divergence

We show that C to T damage curves are strongly influenced by the divergence between the reference genome and the reads. Higher divergence increases mismatch frequencies across all mutation types and reduces the relative C to T damage observed at the end of the reads. This poses a challenge for using C-to-T damage plots to authenticate the ancient origin of microbes without a reference genome. Additionally, cases where the host and microbes are believed to have died simultaneously, similar damage patterns are often interpreted as evidence of their contemporaneity (e.g., (Ferrari et al. 2020; Philips et al. 2017; Feldman et al. 2016; Maixner et al. 2021)). One might thus use the comparison of damage patterns between microbes and their hosts as evidence of coexistence, on the assumption that similar damage profiles imply contemporaneous deposition. Our results show that DNA damage estimates also depend on the phylogenetic closeness of the reference genome. For these reasons, DNA damage patterns can typically only be compared in a relative manner and distinct DNA damage patterns between two samples do not necessarily indicate that they are not contemporaneous. However, our findings suggest that this approach is unreliable when exact reference genomes for the identified microbes are unavailable. Additionally, the biochemical properties of bacterial or viral DNA may affect their preservation differently compared to host DNA, as demonstrated in the case of *Mycobacterium leprae* versus human DNA (Schuenemann et al. 2013).

However, we discovered that damage curve intensity changes in a predictable manner depending on the reference divergence, which could be used to correct the observed damage plots so that they closer match the true values. To achieve this, we first constructed a C-to-T mismatch frequency matrix. This matrix was built to correct read site-specific damage using the simulated damage profiles from references with over 99% identity to the reads . It includes corrected values for all damage types across a range of damage levels (0 -30%) and accounts for the average read identity to the reference genome, with a minimum threshold of 94.7%.

This approach enables some correction to the measured damage signals by first analyzing the observed damage plot and reference divergence percentage. We then identify the closest matching curve within the matrix and apply the calculated site specific corrections (**FigS1**). To evaluate the applicability of this approach to empirical data, we conducted experiments on ancient DNA sequences from various organisms, including *Yersinia pestis, Clostridium tetani, Salmonella enterica*, and *Phytophthora cactorum* (a fungal-like plant pathogen) (**FigS2**). The corrected curves approached the true damage curves, making them more closely aligned to the true values, albeit with some variation. While we acknowledge these corrections can not recapitulate the exact true DNA damage, this approach could serve as a promising starting point for further DNA damage correction algorithms that can consider additional parameters besides reference divergence.

## DISCUSSION

This study provides insights into how reference genome divergence and read depth influence the observed DNA damage patterns in ancient DNA datasets. Under ideal conditions, with a high number of reads mapped to a closely related reference genome, DNA damage plots exhibit the characteristic ‘smiley’ pattern indicative of authentic aDNA. However, such ideal conditions are often not attained in ancient metagenomic studies when working on non-model organisms. Microbial genomic databases often lack comprehensive representation, reducing the likelihood of having access to a closely related reference genome for many ancient microbial taxa. Consequently, researchers working with ancient samples frequently contend with low read counts and divergent reference genomes, which can obscure authentic damage patterns and potentially lead to false-negative conclusions regarding the ancient origin of microbial DNA.

This challenge is even more pronounced in viral aDNA analysis. Viruses are characterized by high mutation rates (Gago et al. 2009), resulting in greater divergence from available reference genomes. Additionally, viral taxa are significantly underrepresented in genomic databases (Kieft and Anantharaman 2022; Lu et al. 2025; Kim, Whon, and Bae 2013). This combination of high sequence divergence and poor reference representation amplifies the sensitivity of damage detection to alignment distance, complicating the interpretation of viral aDNA signals. Our examination of the Hepatitis B virus as a case study further illustrates how reference genome divergence can mask genuine ancient viral DNA damage patterns, highlighting that DNA damage signals should be interpreted in the broader context of the data. Furthermore, while smooth damage curves provide a clearer ancient DNA authentication, such plots generally require extensive sequencing depth and a closely related reference genome (Gago et al. 2009; Der Sarkissian et al. 2021). Our findings demonstrate that even when reads are mapped to a distant organism, a damage signal can be detected if enough reads are available. These results can thus guide researchers in assessing the authenticity of ancient DNA signals under varying data conditions.

## Data availability

All datasets in this paper were either simulated or taken from publicly available datasets which are cited all along the paper. All the code generated for this paper can be found under the github repository : https://github.com/BenjaminGuinet/Impact_ref_on_Ancient_DNA_pattern

## Material and Methods

In this study, we made us of both simulated data, using the *Yersinia pestis* genome as a reference for our read simulations, as well as validating our observations using empirical dataset from the public repositories listed in the SPAAM metadata table (Fellows Yates et al. 2021) accessible at https://www.spaam-community.org/AncientMetagenomeDir/#/.

### Empirical Dataset Analysis

In the first part of this study, we used empirical data from a bacterial and a viral system. For the bacterial dataset, we selected a targeted-capture ancient DNA sequence library (ENA: ERR2862147) from *Yersinia pestis* (GCF_000222975.1). We then mapped these reads against the following reference genomes; *Yersinia bercovieri* (CP124240.1-CP124241.1), and a distant outgroup *Escherichia coli* (U00096.3). Randomly downsamples subsets of the data containing 10,000, 500, and 100 reads respectively were used to assess the effect of read number on DNA damage estimates. For the viral dataset, we used the targeted-capture library (GSA:CRX838871) derived from an ancient Hepatitis B virus (HBV) (Sun et al. 2024). HBV reads were mapped against a standard HBV reference (LC784057.1), two divergent HBV genotypes (from groups H (LC491577.1) and G (AB625343.1)), and the more distantly related Woolly Monkey Hepatitis B virus (AF046996.1). The mapping tools and parameters and the tools used to assess damage patterns were identical to those employed in the simulations described below.

### Simulations and study design

To validate our observations from the empirical dataset, we relied on simulation to systematically explore a range of scenarios and assess their effect on the resulting damage patterns. Specifically, we simulated reads of varying lengths and damage levels, modified the number of reads mapped to the reference genome, and tested different degrees of sequence divergence between the simulated reads and the reference. This approach allowed us to evaluate how each parameter influenced the accuracy and detectability of damage signals.

Our initial simulation involved selecting reads from the reference genome of *Yersinia pestis* (GCF_000222975.1) using NGSNGS v0.9.1 (Henriksen, Zhao, and Korneliussen 2023) **(Fig2-A**). The damage modeling included the following parameters:empirical misincorporation (0% damage, 1% damage (-m b,0.024,0.36,0.03089,0.000366”, labeled as “1DOperc”), 5% damage (-m b,0.024,0.36,0.1545,0.00183”, labeled as “5DOperc”), 10% damage (-m b,0.024,0.36,0.3089,0.00367”, labeled as “10DOperc”), 15% damage (-m b,0.024,0.36,0.4634,0.0055”, labeled as “15DOperc”), 20% damage (-m b,0.024,0.36,0.6179,0.00732”, labeled as “20DOperc”), 25% damage (-m b,0.024,0.36,0.772,0.0091”, labeled as “25DOperc”), and 30% damage (-m b,0.024,0.36,0.926,0.011”, labeled as “30DOperc”). These parameters enabled us to incrementally model DNA damage from 0% to 30%, reflecting varying degrees of cytosine deamination typical in ancient DNA samples. To also capture the fragmentation patterns observed in ancient DNA, we included different read length distributions using the following parameters: 20-49 bp (-ld Uni,20,49”, labeled as “ReadL20-49”), 50-69 bp (-ld Uni,50,69”, labeled as “ReadL50-69”), 70-89 bp (-ld Uni,70,89”, labeled as “ReadL70-89”). Only endogenous DNA was simulated without (modern) contamination.

To model the divergence between the reads and the reference genome, we added divergence directly in the simulated reads (averaging from 94% to 100% identical) using a custom python script (github:https://github.com/BenjaminGuinet/Impact_ref_on_Ancient_DNA_pattern/tree/main) that uses a mutation matrix previously trained on a multiple sequence alignment of 12 different *Yersinia* species. To align the *Yersinia* sequences, we utilized PanACoTA version 1.4.1 (Perrin and Rocha 2021) to detect single-copy orthologous genes among the various species, employing default settings for annotation, a pangenome parameter of -i 0.7 in cluster mode 1, and a corepers parameter of -t 0.3. Subsequently, we used PanACoTA to conduct de novo alignment for each gene family or orthologous gene individually, using default alignment settings. These alignments were then merged to form a comprehensive alignment composed exclusively of core or persistent microbial genes within the genus. We then used this alignment to generate a biologically meaningful matrix to introduce mutations within the reads.

To model the impact of read depth on damage profile, we randomly subsetted our simulated reads from 100 to 1,000,000 from the BAM files using Samtools v1.21 and shuf v8.32 (Danecek et al. 2021). Next, for generating the damage plot information, we employed DamageProfiler v1.1 (Neukamm, Peltzer, and Nieselt 2021) to account for all mutation types.

### Measurement of damage stochasticity

We aimed to quantify the stochasticity in the C-to-T mismatch rate plots by comparing the C-to-T frequency in simulated data with data obtained from mapping up to 1 million reads (**FigS2-A**). This stochasticity is measured by calculating the sum of absolute differences in C-to-T frequencies at each position between the observed damage profile and an expected smooth curve. The expected curve is generated using a large number of reads mapped (1 million) for the same level of divergence from the reference genome. Higher values indicate a greater deviation from the expected curve, which means increasing of the stochasticity.

### Measurement of damage intensity

Another important metric for assessing the damage profile is the intensity of the observed damage, especially at the read edges. To quantify damage intensity, we focus on the C to T mismatch frequencies at the first three positions of the reads, where damage is often most pronounced. To ensure an unbiased comparison between different conditions (when for instance mapping reads to different reference genomes), we calculate the damage intensity as the average C to T mismatch rate at the first three bases and subtracting the average mismatch frequency of all other mismatch types across all bases in the read. In this way we correct for the higher overall background substitution rate when aligning reads to a divergent reference.

### Statistical analysis

To assess the influence of various factors on cytosine-to-thymine (C-to-T) substitution rates, linear regression models were employed using the Python statsmodels package. Two separate models were constructed to evaluate different dependent variables related to DNA damage patterns.

In the first analysis, the relationship between the damage intensity and three predictor variables—DNA damage level (Damage), sequencing read count (nread), and reference genome percent identity (Ref_percID) was examined. A linear model was defined using the formula “damage intensity ∼ Damage + nread + Ref_percID”. This model was fitted using ordinary least squares (OLS) regression through the ols function from the statsmodels.formula.api module. Model fitting was conducted and the statistical significance of each predictor variable was evaluated through the model summary, which provided estimates of regression coefficients, standard errors, t-values, and associated p-values.

In the second analysis, a similar linear regression model was used to investigate factors affecting the damage stochasticity. The formula “Damage stochasticity ∼ Damage + nread + Ref_percID + Read_Length” was specified to include an additional predictor, read length (Read_Length), alongside the previously mentioned variables. Prior to model fitting, the C_to_T_diff column was converted to a numeric format using the pd.to_numeric function with errors=‘coerce’ to handle any non-numeric entries. This conversion ensures data consistency and prevents errors during model fitting. The linear regression was then performed using the ols function, and the model summary was generated to interpret the statistical contributions of each variable.

### Correction of reference bias on damage plot

The tool we developed to apply this correction, named “Correct_damage_ref_bias.py,” takes as input the BAM file (-b) and the matrix (-m), and outputs a plot displaying both the observed and corrected C-to-T damage (**FigS1**). To correct the C-to-T substitution rates induced by postmortem damage in ancient DNA (aDNA) sequences, we developed a two-step computational procedure that accounts for both sequence divergence and damage intensity. All computations were performed in Python using the pandas and numpy libraries. We began by generating a baseline correction matrix using simulated datasets. Specifically, we computed the average corrected C-to-T read depth under low-divergence conditions (Y.pestis simulated reads mapped to Y.pestis reference genomes). We selected records where the number of reads (nread) was equal to 1,000,000 and the average percentage sequence identity (Avrg_Perc_ID) exceeded 99%. These were grouped by damage level and individual C-to-T position along the reads, and the mean value for each group was computed. This provided a reference matrix representing expected damage levels in nearly error-free mapping conditions.

In parallel, we computed the observed mean Corrected_total_C_to_T_DP for each combination of Avrg_Perc_ID and damage level across the entire dataset, under the same read depth (1,000,000). These two datasets (baseline and observed) were merged to allow direct comparison between observed and ideal damage rates across varying divergence levels. This generated a dense matrix estimating the C-to-T profile for any intermediate combination of sequence identity and damage level. To determine the extent of overestimation or underestimation due to mapping divergence, we computed the deviation of each interpolated observed value from the reference (baseline) at high identity. These differences were calculated separately for Observed_C>T0, Observed_C>T1, and Observed_C>T2 and added to the matrix as correction terms . This matrix provides a per-context and per-divergence correction value for adjusting postmortem C-to-T substitution estimates. The final correction matrix was exported as a delimited table for use in downstream post-processing and visualization.

### Supplementary figures

**Fig.S1.**
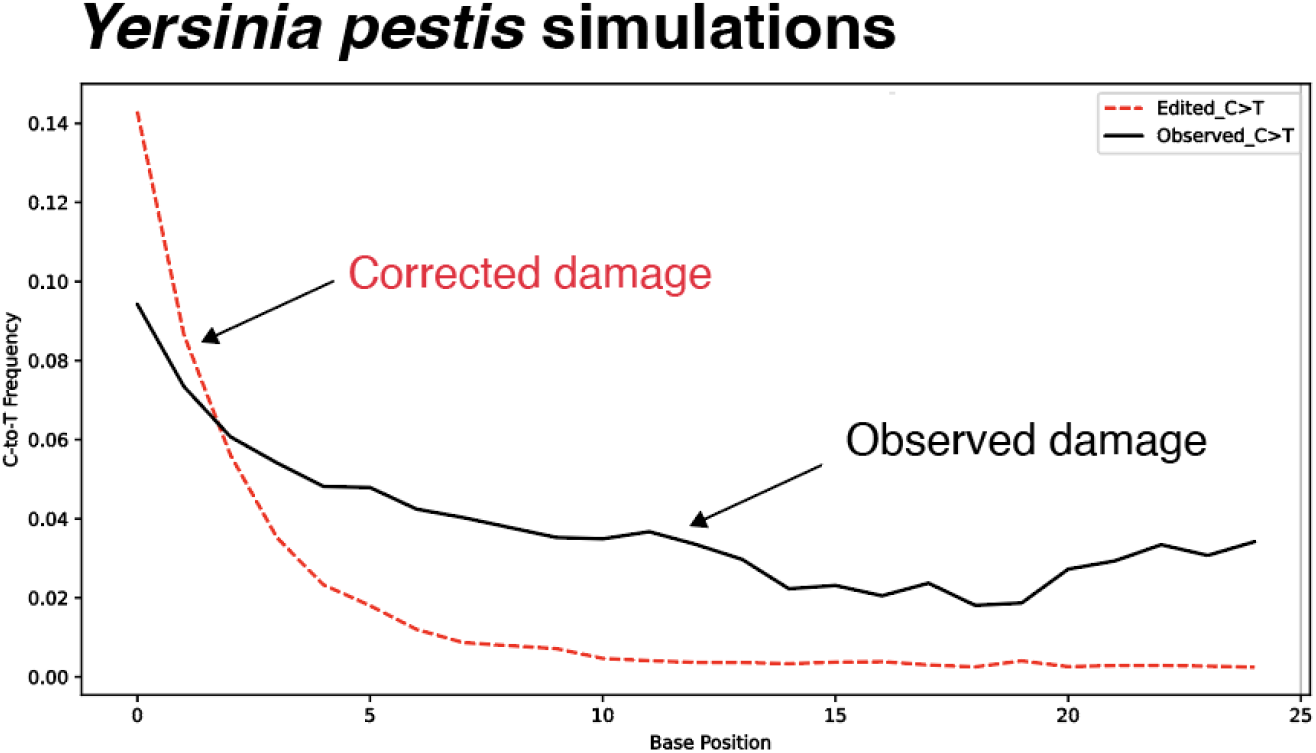
Observed vs. expected C→T damage plot using divergent vs. correct reference genome. As we can observe, the black curve represents the curve obtained when reads are mapped to a divergent reference assembly (93.6% divergence), and the red curve represents the new curve we estimate to be the through if the reads were mapped to the correct reference genome.

**FigS2.**
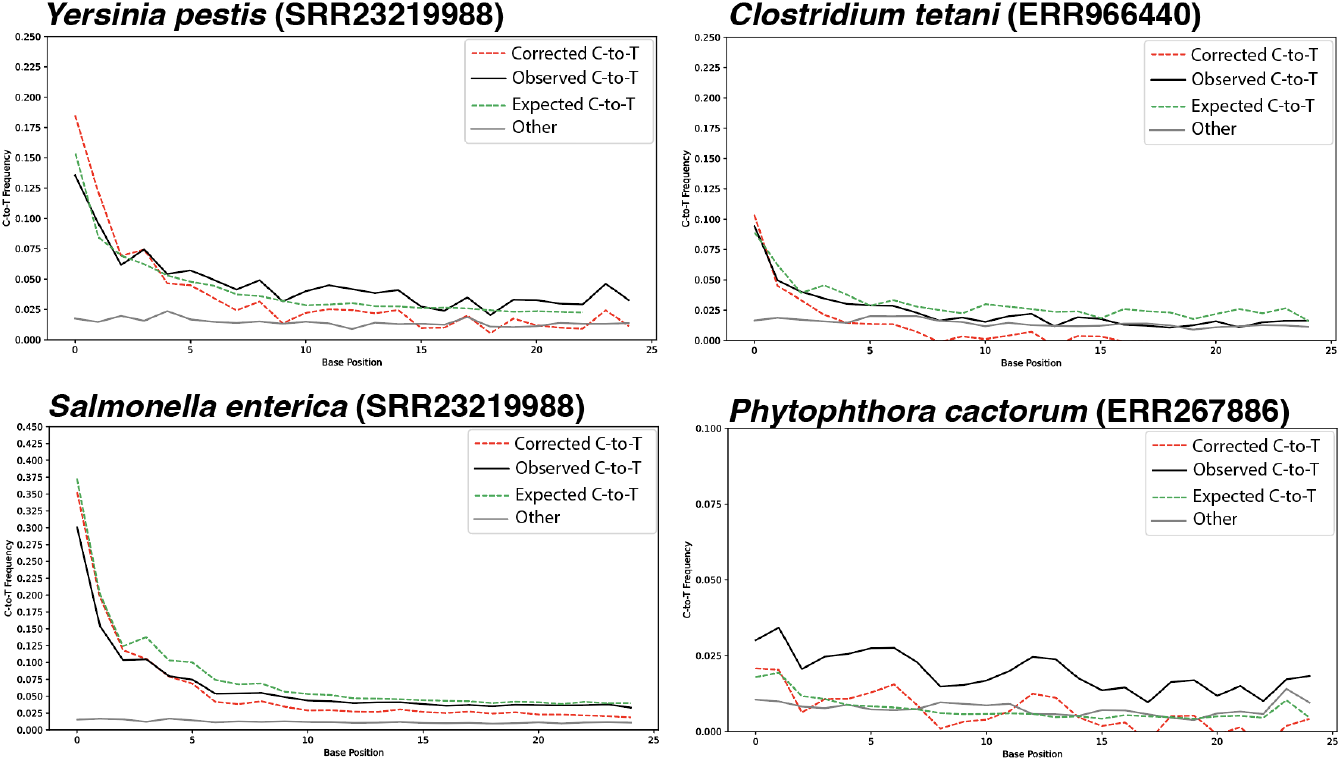
Damage profile before and after reference-bias correction. C→T substitution frequency profiles across the first 25 base positions in ancient DNA samples from *Yersinia pestis* (SRR23219988), *Clostridium tetani* (ERR966440), *Salmonella enterica* (SRR23219988), and *Phytophthora cactorum* (ERR267886). The plots display observed C→T misincorporation rates (black solid lines), corrected C→T frequencies after damage correction (red dashed lines), and the expected C→T frequencies based when reads are mapped to the correct assembly (green dashed lines), and other substitution types (grey solid lines).

**FigS3.**
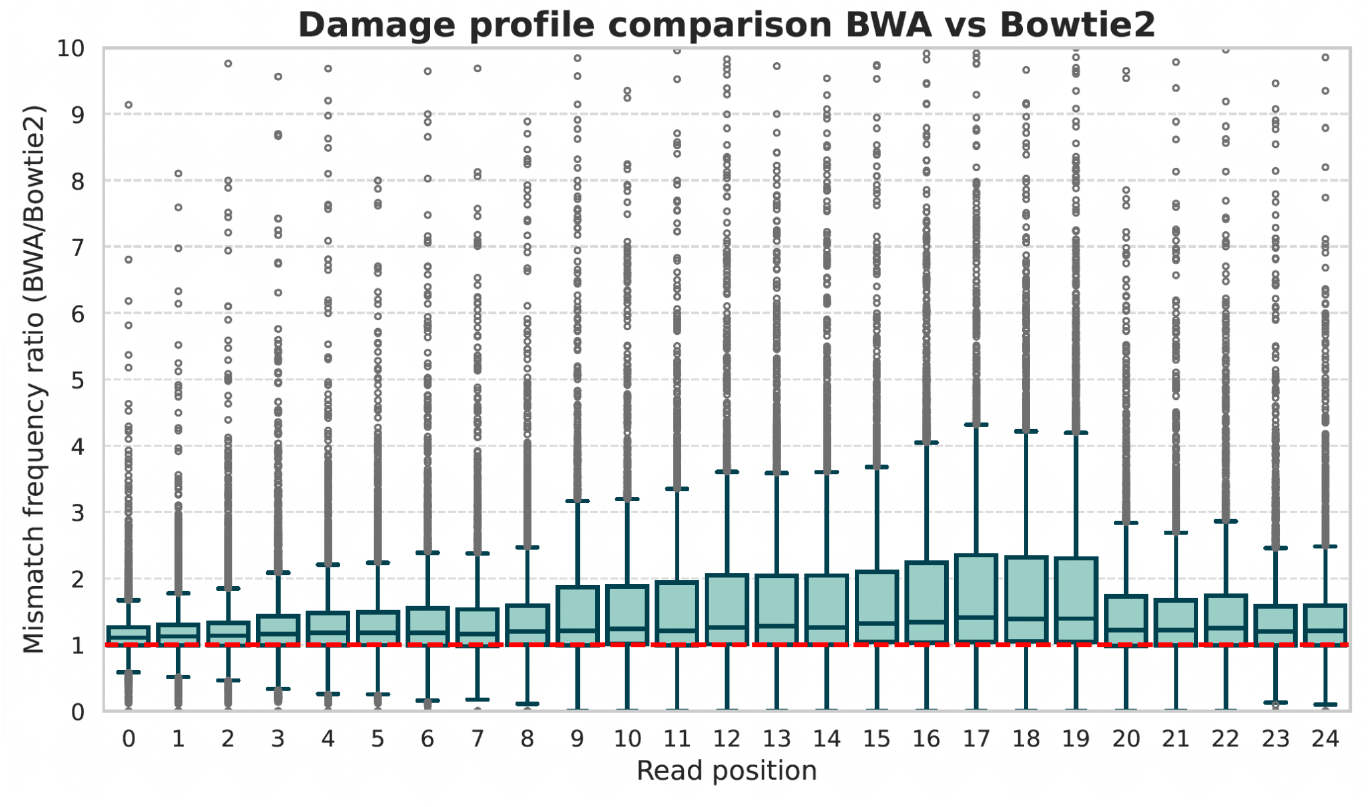
Comparison of damage profiles between BWA and Bowtie2. The Y-axis represents the ratio of mismatch frequency at a given read position (x-axis) in the BWA analysis divided by the corresponding mismatch frequency in Bowtie2. A ratio > 1 indicates that BWA shows a higher mismatch frequency at that position compared to Bowtie2. The red dashed line represents the scenario where the C to T mismatch frequency is identical in both tools. We removed from the plot all simulations that contained only 100 reads to reduce stochasticity.

**FigS4.**
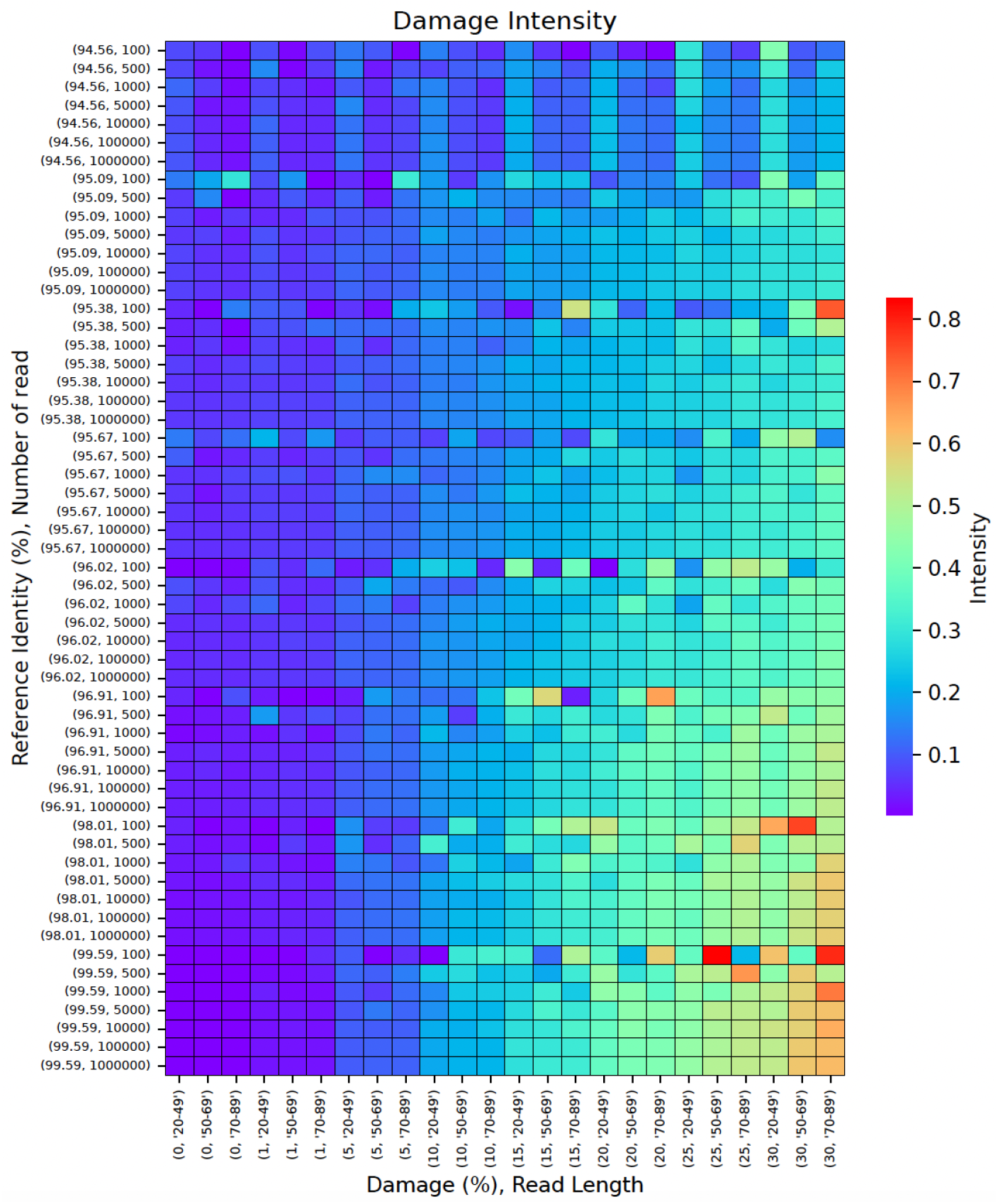
Damage intensity heatmap as a function of damage percentage, read length, and reference identity percentage. The x-axis delineates damage percentages (0% to 30%) across read lengths of 20, 50, and 89 base pairs, while the y-axis represents reference identity percentages ranging from 94.56% to 99.59%, with the number of reads varying from 100 to 1,000,000. The color scale, transitioning from blue to red, indicates the degree of damage intensity, where blue denotes lower damage intensity and red signifies higher damage intensity.

**FigS5.**
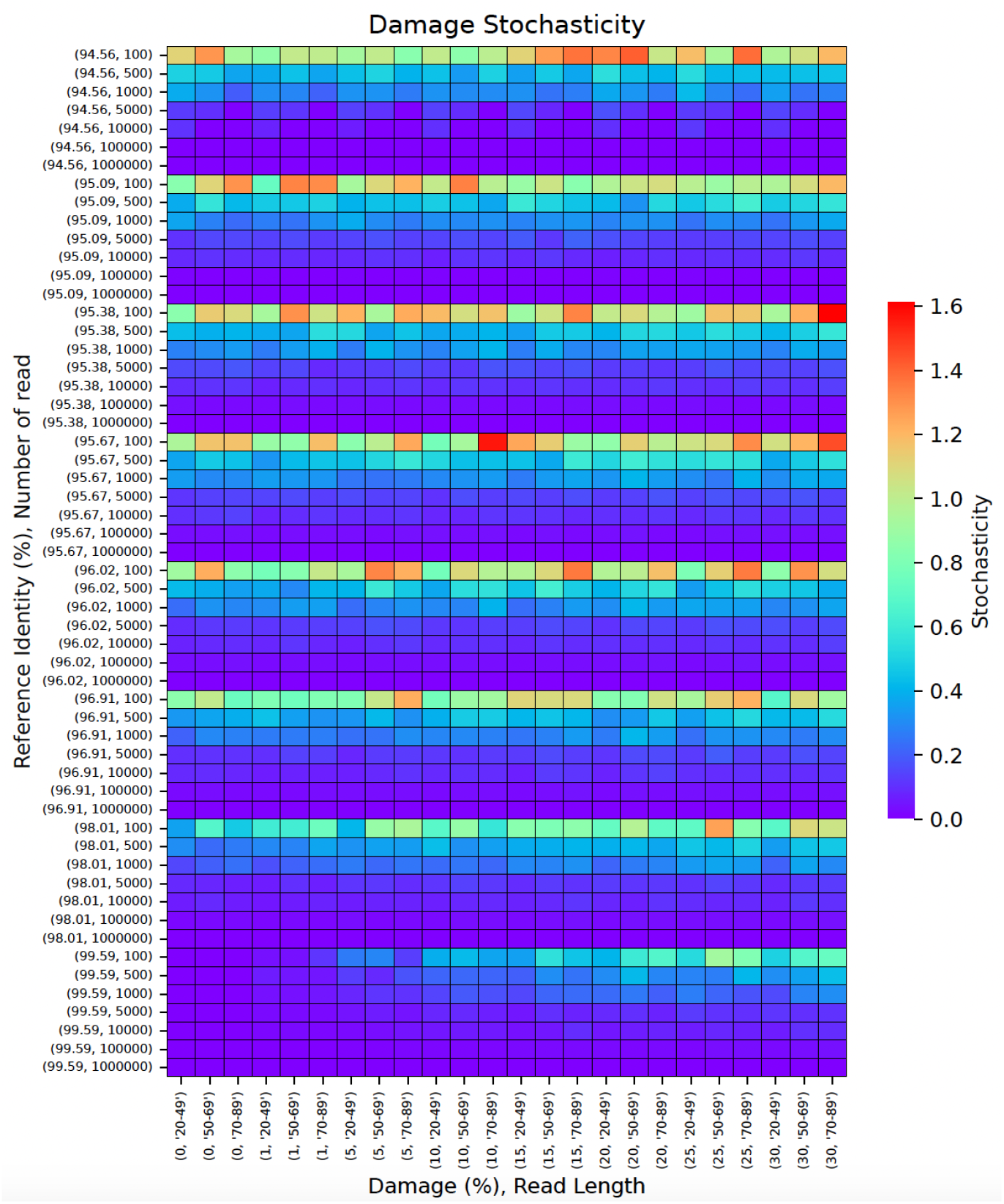
Damage stochasticity heatmap as a function of damage percentage, read length, and reference identity percentage. The x-axis delineates damage percentages (0% to 30%) across read lengths of 20, 50, and 89 base pairs, while the y-axis represents reference identity percentages ranging from 94.56% to 99.59%, with the number of reads varying from 100 to 1,000,000. The color scale, transitioning from blue to red, indicates the degree of damage stochasticity, where blue denotes lower damage stochasticity and red signifies higher damage stochasticity.

**FigS6.**
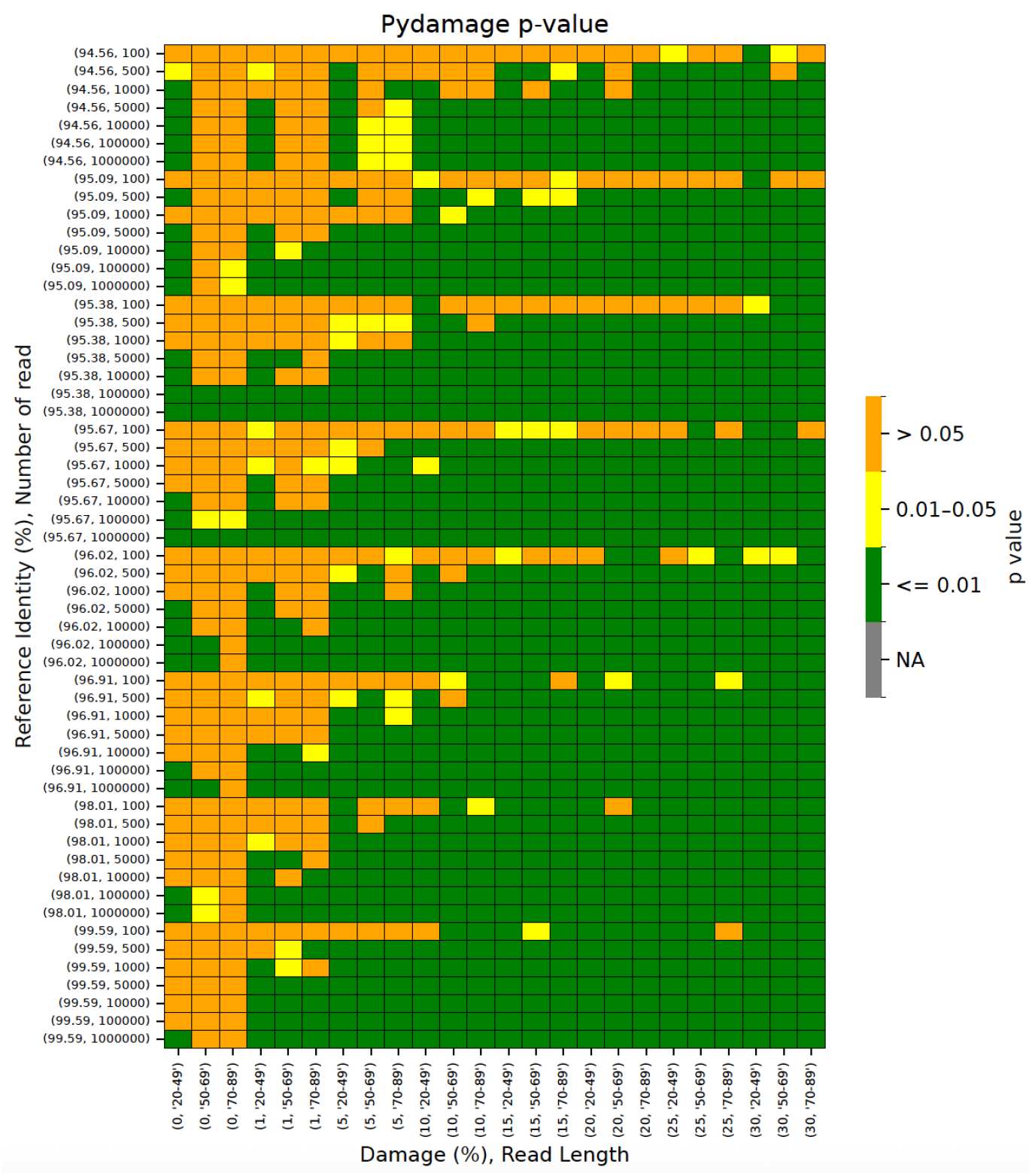
Pydamage pvalues heatmap as a function of damage percentage, read length, and reference identity percentage. The x-axis shows damage percentages ranging from 0% to 30% for read lengths of 20, 50, and 89 base pairs. The y-axis represents reference identity percentages, spanning from 94.56% to 99.59%, with the number of reads varying between 100 and 1,000,000. The color scale transitions from orange to green, indicating the significance of p-damage. Orange signifies a high p-value, suggesting no significant damage, while green indicates a low p-value, denoting significant damage.

